# Connectome-based predictive modeling of empathy in adolescents with and without the low-prosocial emotion specifier

**DOI:** 10.1101/2022.10.14.512331

**Authors:** Drew E. Winters, Anika Guha, Joseph T. Sakai

**Affiliations:** Department of Psychiatry, University of Colorado School of Medicine, Anschutz Medical Campus

**Author notes:** Corresponding author: Drew E. Winters.

**Keywords:** Connectome-based predictive modeling, functional connectivity, empathy, callous-unemotional traits, adolescents

## Abstract

Empathy impairments are an important part of a broader affective impairments defining the youth antisocial phenotype callous-unemotional (CU) traits and the DSM-5 low prosocial emotion (LPE) specifier. While functional connectivity underlying empathy and CU traits have been well studied, less is known about what functional connections underly differences in empathy amongst adolescents qualifying for the LPE specifier. Such information can provide mechanistic distinctions for this clinically relevant specifier. The present study uses connectome-based predictive modeling that uses whole-brain resting-state functional connectivity data to predict cognitive and affective empathy for those meeting the LPE specifier (n= 29) and those that do not (n= 57). Additionally, we tested if models of empathy generalized between groups as well as density differences for each model of empathy between groups. Results indicate the LPE group had lower cognitive and affective empathy as well as higher CU traits and conduct problems. Negative and positive models were identified for affective empathy for both groups, but only the negative model for the LPE and positive model for the normative group reliably predicted cognitive empathy. Models predicting empathy did not generalize between groups. Density differences within the default mode, salience, executive control, limbic, and cerebellar networks were found as well as between the executive control, salience, and default mode networks. And, importantly, connections between the executive control and default mode networks characterized empathy differences the LPE group such that more positive connections characterized cognitive differences and less negative connections characterized affective differences. These findings indicate neural differences in empathy for those meeting LPE criteria that may explain decrements in empathy amongst these youth. These findings support theoretical accounts of empathy decrements in the LPE clinical specifier and extend them to identify specific circuits accounting for variation in empathy impairments. The identified negative models help understand what connections inhibit empathy whereas the positive models reveal what brain patterns are being used to support empathy in those with the LPE specifier. LPE differences from the normative group and could be an appropriate biomarker for predicting CU trait severity. Replication and validation using other large datasets are important next steps.

---

Callous-unemotional (CU) traits is a youth antisocial phenotype involving profound affective impairments in prosocial emotions of guilt, remorse, and empathy [1, 2]. The presence of CU traits has become an important specifier in the clinical assessment of psychiatric disorders including conduct and oppositional defiant disorder known as the low prosocial emotion (LPE) specifier [DSM-5: 3, ICD-10: 4]. While these disorders associate with aggression and violence that impact society [2, 5] the presence of the LPE specifier indicates youth with more severe, stable, and chronic antisocial behavior [5, 6].

Low empathy is an important impairment amongst a broader set of affective and interpersonal impairments underlying the LPE specifier [1, 2, 7]. Where empathy is broadly defined as the capacity to identify and resonate with another’s affective state [8], the cognitive component (i.e., cognitive empathy, aka: perspective taking or theory of mind) involves the cognitive inference of another’s affective state whereas the affective component (i.e., affective empathy) is the affective resonance with what another is feeling [9]. These prosocial emotions are a precursor to prosocial behavior [8, 10]; however, their absence (e.g., in the LPE specifier) is thought to underlie propensity for substance use [11, 12] and antisocial behavior [1, 2].

Given the LPE specifier involves lower levels of empathy, it is plausible that these differences would be reflected in the brain. Contemporary brain research demonstrates that, as opposed to the activation of a specific brain region, behaviors are subserved by the interactions between multiple regions of the brain, which can be measured using functional connectivity [13]. Functional connectivity studies demonstrate that cognitive empathy positively associates with connectivity in the default mode network [DMN; 14, 15] as well as executive control network [ECN; 16]; however, CU traits are associated with less connectivity in the DMN [17] in relation to cognitive empathy [15, 18] and less positive connectivity in the ECN [19]. Affective empathy associates with greater connectivity in the salience network [SAL; 20, 21] and less connectivity between the SAL and DMN [15]; however, CU traits associates with less connectivity in the SAL [22] and more connectivity between the SAL and DMN [15]. General support of both cognitive and affective empathy involves a competition between the ECN and DMN meaning we expect fewer positive connections between these networks in relation to empathy [23] but CU traits associates with more positive connections between ECN-DMN [19] and CU traits moderates the association between ECN-DMN connectivity in relation to empathy [24].

Importantly, this prior work focuses on CU traits as a dimensional construct when there may be clinically relevant distinctions between those meeting DSM-5 LPE criteria versus those that do not. Moreover, although these studies demonstrate differences in functional connectivity amongst the brains of those higher in CU traits – they focus exclusively on the three major networks (DMN, ECN, and SAL), though other networks are likely involved. For example, affective empathy also involves sensory motor (SMN) and Limbic networks [25]. And, both cognitive and affective empathy are more generally supported by the occipital [26] and cerebellar networks [27]. It is plausible that there are broader patterns in the functional connectomes of those qualifying for the LPE specifier representing deficits in cognitive and affective empathy.

The current study examined differences in empathy across the entire functional connectome amongst adolescents with and without the LPE specifier. Specifically, we used connectome predictive modeling [28, 29], a widely applied approach for predicting human behavior and traits that has been used to predict CU traits [30] and empathy in clinical samples [31]. This approach was used to identify positive and negative models of functional connections underlying cognitive and affective empathy amongst those with and without the LPE specifier. We hypothesized that distinct models of brain connections predicting cognitive and affective empathy for those with and without the LPE specifier would be identified and that the model predicting empathy would not generalize between these groups. Moreover, consistent with prior research on density in CU traits, we hypothesized there would be differences in network density – particularly in the DMN, ECN, SAL and between these networks. Such information will help identify mechanistic differences underlying empathy between clinically meaningful distinctions of CU traits.

## Methods

### Preregistration and Transparency

The present analysis was preregistered as a secondary data analysis (https://osf.io/svzg4). While the primary aims and analyses remained the same, during our analyses we revealed substantial differences between those with and without the LPE specifier and we added analyses characterizing these differences by examining the density of connections.

### Sample

Participants raw fMRI data were downloaded from the Rockland study collected by the Nathan Kline Institute via the 1000 connectomes project (www.nitrc.org/projects/fcon_1000/). Participants in adolescence (13-17 years-old) with an IQ > 80 as measured by the WAIS-II [α = .96; 32] were included. We excluded potential participants with motion > 3mm or > 20% of invalid fMRI scans. Two participants had spikes in motion that was near the end of the session that were able to be retained by cutting their time series (still retaining > 90% of the time series) leaving a total analysis sample of 86. These participants were predominantly White (White= 63%, Black = 24%, Asian = 9%, Indian = 1%, other= 3%), with a mean age of 14.5 (14.52±1.31) years, and slightly more males (females = 48%). Consent for all participants as well as data collection procedures are outlined in Nooner, Colcombe [33].

### Measures

### Inventory of Callous-Unemotional Traits (ICU)

CU traits were assessed using the ICU total score (Frick, 2004), which was used to calculate the low prosocial emotion (LPE) specifier according to recommended standards by Kimonis, Fanti [6]. There are four ways to compute the specifier using either four or nine items as well as split and extreme coding methods. We Identified anyone qualifying for any of the specifiers as qualifying for LPE specifier group and those not qualifying for any calculations as the normative group.

### Conduct Problems

Conduct problems were assessed using the raw scores of the externalizing subscale of the Youth Self Report [34]. Items are rated on the three-point scale (0 not true – 2 very true) indicating level of conduct problems expressed over the previous 6 months. This scale had adequate reliability in the current sample (α=0.87).

### Tanner Stage

The Tanner assessment was used to measure sex and pubertal stage (α = 0.77). Parents of participants rated pictures of secondary sex characteristics on a scale of 1 (pre-pubertal) to 5 (full maturity) indicating pubertal development [35].

### fMRI Acquisition

Resting state fMRI images were collected by the Nathan Kline Institute using a Siemens TimTrio 3T scanner with a blood oxygen level dependent (BOLD) contrast and an interleaved multiband echo planar imaging (EPI) sequence. Each scan involved resting state (260 EPI volumes; repetition time (TR) 1400ms; echo time (TE) 30ms; flip angle 65°; 64 slices, Field of view (FOV) = 224mm, voxel size 2mm isotropic, duration = 10 minutes) and a magnetization prepared rapid gradient echo (MPRAGE) anatomical image (TR= 1900ms, flip angle 9°, 176 slices, FOV= 250mm, voxel size= 1mm isotropic). Removing scans was not necessary because the Siemens sequence does not collect images until T1 stabilization is achieved.

### Resting-State fMRI Preprocessing

Imaging data was preprocessed with the standard preprocessing in the CONN toolbox [version 18b; 36] that uses Statistical Parametric Mapping [SPM version 12; 37]. Motion outliers were flagged for correction if > 0.5mm using the Artifact Detection Tools (ART; http://www.nitrc.org/projects/artifact_detect) and regressed out using spike regression. Because of the multiband acquisition, slice timing correction was not used [38, 39]. The anatomic component-based noise correction [aCompCor; 36] was used to regress out white matter and CSF noise. MPRAGE and EPI images were co-registered and normalized to an MNI template. The data was bandpass filtered to retain resting state signals between 0.008 and .09HZ. Finally, this data parcellated into 164 ROIs using the Harvard Oxford atlas for cortical and sub-cortical areas [40] as well as the Automated Anatomical Labeling Atlas for cerebellar areas [41]. Regions were assigned to one of seven networks: Default Mode (DMN), Executive Control (ECN), Salience (SAL), Limbic, Occipital, and Cerebellar.

A total of 24 participants were found to have excess motion > 3mm and four had >20% of invalid scans. We were, however, able to retain two of the participants with excess motion that was at the end of the timeseries, which we were able to snip it while still retaining > 90% of the timeseries. This left a total of 86 participants for analysis.

### Individual-Level Functional Connectomes

Individual-level resting state connectomes were derived as correlation matrices using the ConnectivityMeasure command from the package “NiLearn” [42] across all 164 parcellated nodes. Time courses for each individual were entered individually, which resulted in individual-level connectomes.

## Analysis

### Connectome based predictive modeling (CPM)

CPM [28, 29] was conducted to predict individual differences in cognitive and affective empathy based on whole brain resting state functional connectivity using python (https://github.com/YaleMRRC/CPM) scripts and code used for the present study available at https://github.com/drewwint/pub_CPM_emp_LPE and OSF project at https://doi.org/10.17605/OSF.IO/NME25. First the data was separated by those with and without the LPE specifier. Following CPM procedures, we the conducted feature selection to identify positive and negative models for each group in relation to cognitive and affective empathy. Because p-value thresholds are arbitrary, we tested thresholds ranging from 0.005 to 0.01 [e.g., 43], which revealed the best fitting models at the threshold of p= 0.005. Next, these identified features for each group and outcome were used for predicting cognitive and affective empathy using machine learning and permutation tests of this model was used to determine significance.

### Elastic net regression, cross-validation, and evaluation

For cognitive and affective empathy predictions for positive and negative connectomes, a linear elastic net regression was implemented in the python package “scikit-learn” [44] to evaluate if information about connectomes of those with and without LPE specifier accounts for variance in cognitive and affective empathy. Model generalization was evaluated using a nested five-fold cross-validation procedure involving hyper-parameter tuning and cross-validation. We used mean squared error, R^2^, and mean absolute error to evaluate the model as well as comparing training and testing cross-validation scores. The model was assessed comparing mean squared errors from each fold to a dummy model as well as a permutation test on R^2^ with 2000 iterations. Results were considered statistically significant if 95% of these 2000 R^2^ values were lower than the R^2^ of the real data [45]. For all analyses, we controlled for conduct problems, sex, Tanner stage, and head motion.

### Network Analyses

Next, we examined network contributions to prediction of cognitive and affective empathy for each group and each model. Density of connections within and between network were derived from individual connectomes and aggregated for each predictive model (positive and negative cognitive and affective empathy by LPE/normative group). Heat maps were created for each of these, and t-tests were conducted for within and between network connections to determine differences in density between theoretically derived LPE groups. For between network connections, consistent with the majority of research on between network connections and CU traits [e.g., 15, 19, 46], we focused on density between the DMN, ECN, and SAL networks.

## Results

Adolescents in the normative group were significantly higher on cognitive and affective empathy (t= 3.52, p_perm_ < 0.001) while also lower in both conduct problems (t= 7.06, p_perm_ < 0.001) and CU traits (t= −9.44, p_perm_ < 0.001; Table 1). There were more participants that were White in the normative group relative to the LPE group (Table1), thus in subsequent analysis we ensured we included race as a covariate to account for spurious effects related to this difference in distribution. Sex did not significantly associate with any other variables in the analysis, race associated with age and tanner (Table 1). As expected, CU traits negatively associated with both cognitive and affective empathy (Table 1).

**Table 1.** Demographics and correlations.

| | | Correlations | | | | | | | Whole | Normative | LPE | t-test or $\chi^2$ | |
| --- | --- | --- | --- | --- | --- | --- | --- | --- | --- | --- | --- | --- | --- |
| | | 2 | 3 | 4 | 5 | 6 | 7 | 8 | M±SD or n(%) | | | Stat | P<sub>perm</sub><sup>b</sup> |
| 1 | Sex<sup>a</sup> | -0.012 | -0.031 | 0.176 | -0.197 | -0.127 | 0.051 | 0.026 |  |  |  |  |  |
|  | Male |  |  |  |  |  |  |  | 45(52%) | 29(34%) | 16(19%) | 0.02 | 0.89 |
|  | Female |  |  |  |  |  |  |  | 41(48%) | 28(32%) | 13(15%) | 0.31 | 0.57 |
| 2 | Race<sup>a</sup> |  | 0.304* | 0.258* | 0.010 | 0.121 | -0.131 | 0.101 |  |  |  |  |  |
|  | White |  |  |  |  |  |  |  | 54(63%) | 38(44%) | 16(19%) | 6.30* | 0.01 |
|  | Non-White |  |  |  |  |  |  |  | 32(37%) | 19(22%) | 13(15%) | 0.31 | 0.58 |
| 3 | Age | | | 0.452* | -0.069 | -0.039 | 0.047 | 0.072 | 14.66±1.36 | 14.7±1.43 | 14.6±1.23 | 0.36 | 0.74 |
| 4 | Tanner | | | | -0.009 | -0.135 | 0.132 | 0.216* | 4.10±0.97 | 4.01±0.99 | 4.25±0.92 | -1.17 | 0.30 |
| 5 | Cognitive | | | | | 0.462* | -0.472* | -0.056 | 12.84±5.01 | 14.1±4.71 | 10.3±4.71 | 3.52* | <0.001 |
| 6 | Affective | | | | | | -0.740* | -0.141 | 17.54±5.10 | 19.8±3.98 | 13.2±4.22 | 7.06* | <0.001 |
| 7 | CU traits | | | | | | | 0.169 | 22.31±9.20 | 17.5±5.77 | 31.5±7.61 | -9.44* | <0.001 |
| 8 | Conduct | | | | | | | | 7.22±5.54 | 6.75±4.63 | 8.14±7.01 | -1.09 | 0.27 |

<sup>a</sup> spearman correlation<sup>b</sup> p-values via 10,000 permutations

### Performance of CPM Predicting Empathy Scores

For cognitive empathy in the LPE group, the positive model did not reliably predict cognitive empathy, however the negative model did (R^2^= 0.63, P_perm_ < 0.001; Table 2, Figure 1). In the normative group the positive model reliably predicted empathy, however the negative model did not (R^2^= 0.52, P_perm_ < 0.001; Table 2, Figure 2). For affective empathy, both positive (R^2^= 0.48, P_perm_ < 0.001) and negative (R^2^= 0.57, P_perm_ < 0.001) models in the LPE group reliably predicted affective empathy. Similarly, in the normative group both positive (R^2^= 0.49, P_perm_ < 0.001) and negative (R^2^= 0.55, P_perm_ < 0.001) models reliably predicted affective empathy (Table 2, Figure 1 and 2).

**Table 2.** Model fits

| Model | Train |  |  | Test |  |  | Permutation |  | Decision |
| --- | --- | --- | --- | --- | --- | --- | --- | --- | --- |
|  | MSE | MAE | R<sup>2</sup> | MSE | MAE | R<sup>2</sup> | R<sup>2</sup> | P<sub>perm</sub> |  |
| LPE |  |  |  |  |  |  |  |  |  |
| Cognitive – positive | 1.37±1.52 | 0.81±0.49 | 0.94±0.07 | 9.06±3.19 | 2.64±0.56 | 0.16±0.66 | 0.19 | 0.05 | Not generalized |
| Cognitive – negative | 0.68±0.51 | 0.60±0.24 | 0.96±0.03 | 4.08±2.65 | 1.66±0.50 | 0.64±0.20 | 0.63 | <0.001 | Good |
| Affective – positive | 0.71±0.39 | 0.66±0.21 | 0.95±0.02 | 5.19±3.34 | 1.80±0.78 | 0.52±0.32 | 0.48 | <0.001 | Good |
| Affective - negative | 0.59±0.38 | 0.59±0.23 | 0.97±0.02 | 4.20±1.12 | 1.71±0.23 | 0.52±0.19 | 0.57 | <0.001 | Good |
| Normative |  |  |  |  |  |  |  |  |  |
| Cognitive – positive | 5.28±0.77 | 1.77±0.12 | 0.76±0.03 | 10.13±4.65 | 2.51±0.67 | 0.48±0.23 | 0.52 | <0.001 | Good |
| Cognitive – negative | 17.11±4.3 | 3.08±0.43 | 0.22±0.15 | 21.8±9.49 | 3.61±1.03 | -0.1±0.26 | 0.02 | 0.03 | Not generalized |
| Affective – positive | 3.51±0.37 | 1.46±0.11 | 0.772±0.02 | 7.06±1.68 | 2.09±0.23 | 0.48±0.11 | 0.49 | <0.001 | Good |
| Affective - negative | 1.77±0.65 | 1.04±0.19 | 0.87±0.04 | 7.12±4.23 | 2.25±0.72 | 0.42±0.28 | 0.55 | <0.001 | Good |

**Figure 1.**
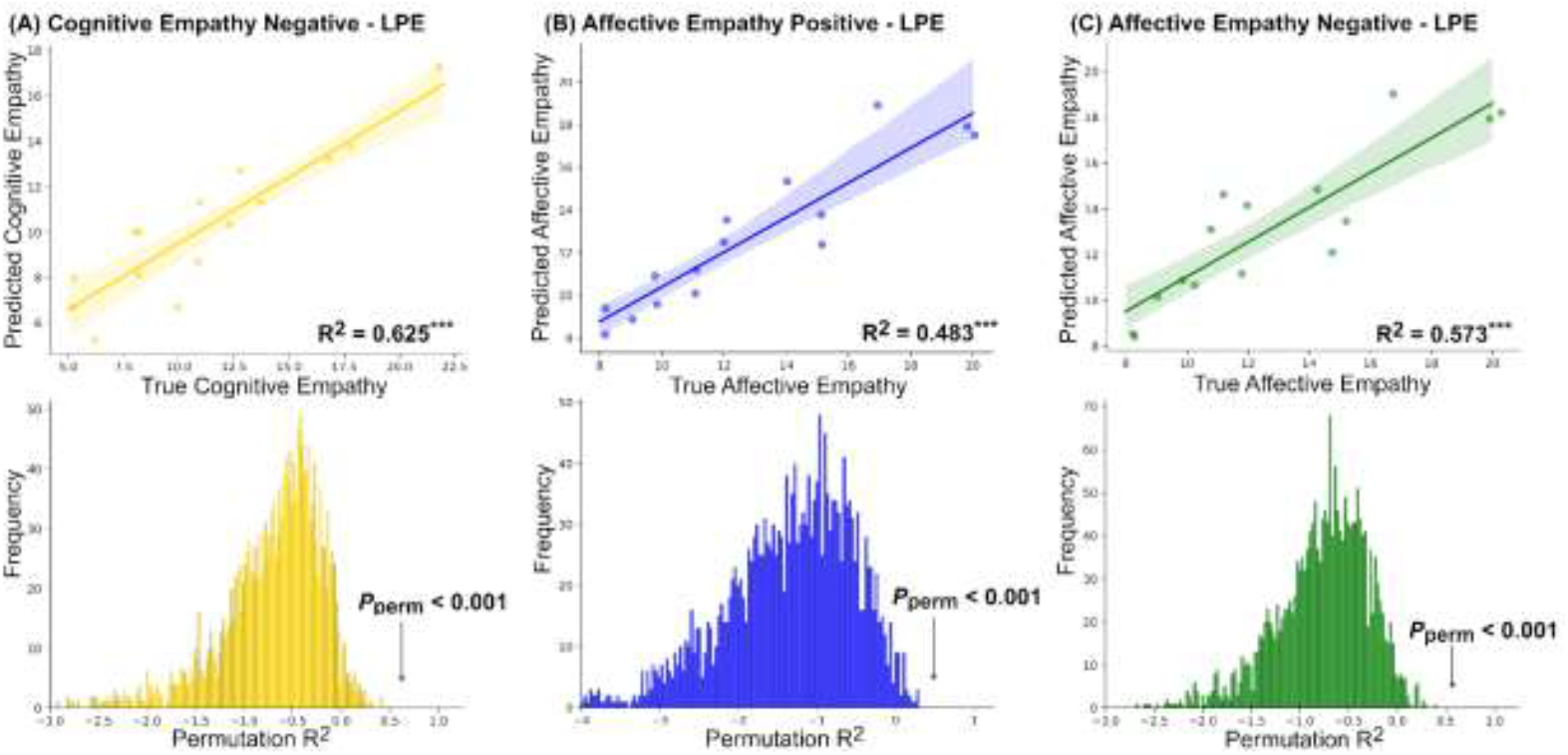
For the low prosocial emotion (LPE) group – results of the negative and positive models of network connectivity predicting cognitive and affective empathy. (A) Negative model predicting cognitive empathy, (B) positive model predicting affective empathy, (C) negative model predicting affective empathy. Note: the positive cognitive model is not pictured because it did not reliably predict cognitive empathy in the LPE group.

**Figure 2.**
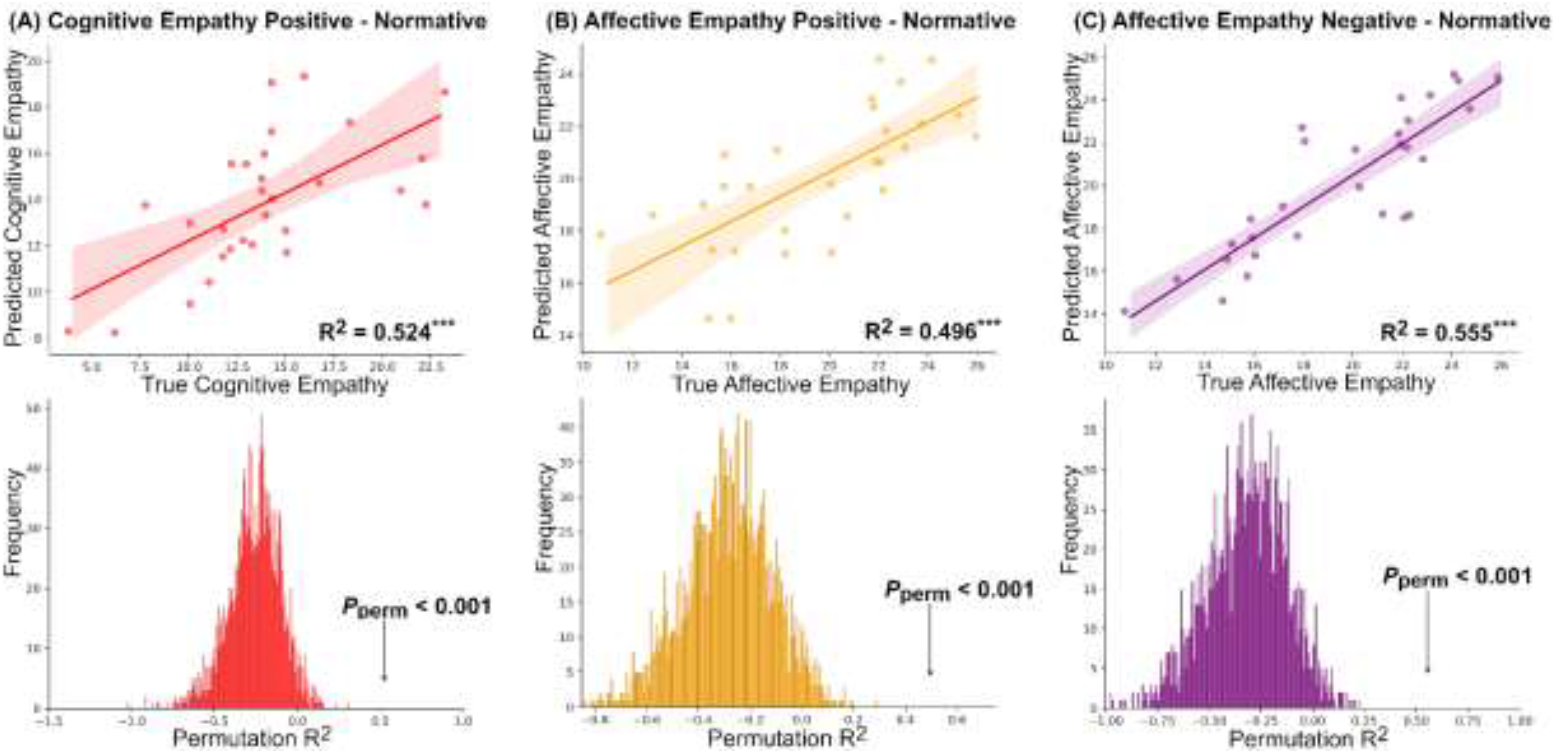
For the normative group – results of the negative and positive models of network connectivity predicting cognitive and affective empathy. (A) Positive model predicting cognitive empathy, (B) positive model predicting affective empathy, (C) negative model predicting affective empathy. Note: the negative cognitive model is not pictured because it did not reliably predict cognitive empathy in the normative group.

### Generalization of Normative Empathy Models to LPE

None of the models, neither positive or negative, from the normative participants generalized to the LPE participants when predicting cognitive or affective empathy (Table 3).

**Table 3.** Generalizing from normative to LPE

|  | R<sup>2</sup><sub>perm</sub> | P<sub>perm</sub> | Decision |
| --- | --- | --- | --- |
| Cognitive positive | -0.610 | 0.889 | Not generalized |
| Cognitive negative | -0.165 | 0.185 | Not generalized |
| Affective positive | -0.225 | 0.972 | Not generalized |
| Affective negative | -0.611 | 0.889 | Not generalized |

### Network Density for Models Predicting Cognitive and Affective Empathy

Figure 3 depicts network densities accounting for variance in cognitive and affective empathy for normative and LPE groups. For cognitive empathy, the LPE negative model had the most connections in the SAL and Occipital networks, as well as moderate connections between the DMN, ECN and SAL, associated negatively with cognitive empathy. The normative positive model had more dense connections in the DMN, ECN, SAL and cerebellar networks that associated positively with cognitive empathy. For affective empathy, the LPE positive model had more dense connections in the ECN and SMN positively associating with affective empathy, whereas the negative model had the most connections within the occipital negatively associating with affective empathy. The normative positive model had the most connections within the SAL positively associating with affective empathy, whereas the normative negative model had the most connections within the ECN and cerebellar negatively associations with affective empathy.

**Figure 3.**
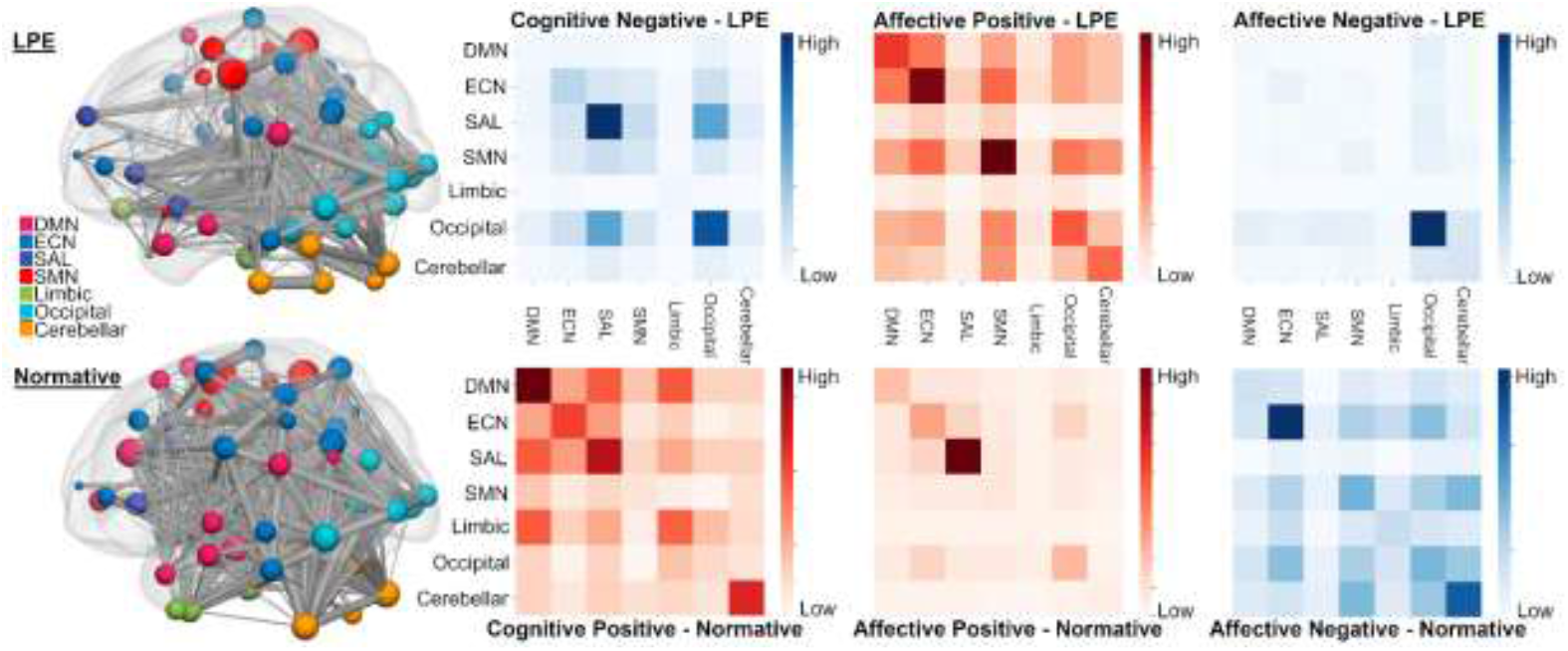
Network density matrices for positive (red) and negative (blue) models of cognitive and affective empathy by low prosocial emotion (LPE) and normative groups.

### Density Differences in Networks Predicting Empathy in Normative and LPE

T-tests revealed density differences between normative and LPE groups for positive and negative models associating with cognitive and affective empathy (Table 4, Figure 4). For positive connections with cognitive empathy, the LPE group had fewer positive connections within the DMN, SAL, SMN, Limbic, and Cerebellar and between SAL-DMN and SAL-ECN as well as more density in the ECN and between ECN-DMN (Table 4, Figure 4). For negative connections with cognitive empathy, the LPE group had less density in the DMN and cerebellar networks and more density in the Occipital and between SAL-DMN. For positive connections with affective empathy, the LPE group had fewer positive connections in the DMN, SAL, and Limbic and between DAL-DMN and SAL-ECN. For negative connection with affective empathy, the LPE group had fewer negative connections in the DMN, ECN, SMN, Limbic, cerebellar and between ECN-DMN (Table 4, Figure 4).

**Table 4.** Network density differences between LPE and normative by network connections

|  | Cognitive |  |  |  | Affective |  |  |  |
| --- | --- | --- | --- | --- | --- | --- | --- | --- |
|  | Positive |  | Negative |  | Positive |  | Negative |  |
|  | T | P<sub>perm</sub> | T | P<sub>perm</sub> | T | P<sub>perm</sub> | T | P<sub>perm</sub> |
| **Within network** |  |  |  |  |  |  |  |  |
| DMN | -5.456* | 0.000 | -8.294* | 0.008 | -2.404* | 0.023 | -5.317* | 0.001 |
| ECN | 2.501* | 0.013 | -0.768 | 0.450 | -0.237 | 0.815 | -7.721* | 0.000 |
| SAL | -10.905* | 0.000 | 1.341 | 0.294 | -6.541* | 0.001 | -1.754 | 0.143 |
| SMN | -4.181* | 0.000 | 0.112 | 0.913 | 1.485 | 0.149 | -7.409* | 0.000 |
| Limbic | -9.022* | 0.000 | 0.488 | 0.714 | -15.336* | 0.067 | -7.681* | 0.006 |
| Occipital | 0.013 | 0.989 | 2.524* | 0.001 | -1.515 | 0.151 | 3.935* | 0.001 |
| Cerebellar | -6.667* | 0.000 | -3.315* | 0.008 | 1.289 | 0.232 | -7.638* | 0.000 |
| **Between network** |  |  |  |  |  |  |  |  |
| ECN-DMN | 3.024* | 0.006 | 0.865 | 0.460 | 1.504 | 0.148 | -4.790* | 0.002 |
| SAL-DMN | -10.895* | 0.000 | 3.283* | 0.002 | -3.550* | 0.001 | -0.437 | 0.667 |
| SAL-ECN | -10.121* | 0.000 | 1.444 | 0.190 | -3.889* | 0.001 | 1.428 | 0.179 |

Note: negative values mean LPE is less dense than normative and positive values mean LPE has more density than normative

**Figure 4.**
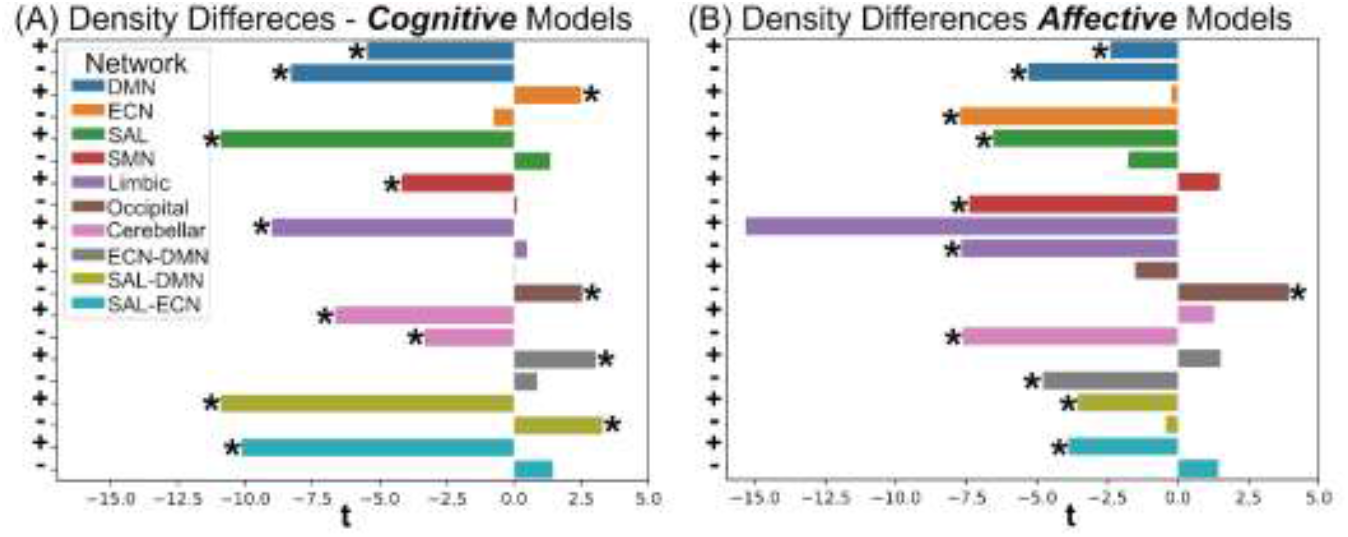
Mean differences in network density between normative and low prosocial emotion specifier (LPE) groups by network. Note: positive network connections noted by (+) are the first for each network and negative network connections noted b (-) are second for each network. (A) Density differences for cognitive empathy models (B) Density differences for affective empathy models.

## Discussion

Amongst a sample of community adolescents, the present results revealed cognitive and affective empathy was lower among adolescents in the LPE group whereas CU traits and conduct problems were higher. Moreover, distinct patterns of connectivity reliably predicted cognitive and affective empathy; however, the models that predicted empathy in those without the LPE specifier did not generalize to those with the LPE specifier. Density differences within and between networks between groups support existing literature but extend prior findings by demonstrating these density differences in relation to cognitive and affective empathy for the clinically relevant LPE specifier explicitly.

### Distinct Patterns of Functional Connectivity Predicted Cognitive and Affective Empathy

Results revealed positive and negative models that predicted cognitive and affective empathy for each group. However, two models did not reliably predict, which included the positive model for cognitive empathy in the LPE specifier group and the negative cognitive empathy model in the normative group. The lack of reliable prediction suggests that a model of brain connections positively predicting cognitive empathy did not exist for the LPE specifier group – meaning that there is a greater likelihood for the patterns of functional connectivity in those with the LPE specifier to negatively associate with cognitive empathy. Likewise, for the normative group, functional connectivity is more likely to associate positively, as opposed to negatively, with cognitive empathy.

Affective empathy models found both positive and negative models for both groups that demonstrated different patterns. This suggests that distinct patterns of functional connectivity can support and also oppose affective empathy for both groups. Importantly, the patterns observed for each of these groups are different.

### Models Predicting Empathy Did Not Generalize Between Low Prosocial Emotion Groups

Connectivity patterns found for both positive and negative models in the normative group did not generalize to the LPE specifier group. One interpretation of these findings is that there are inherently different patterns of functional connections explaining cognitive and affective empathy between normative and clinically relevant LPE specifiers. This fits the overall theoretical understanding that there are substantial differences amongst those higher in CU traits and qualify for the LPE specifier having differences in empathy function. Here we extend the support for this view by evidencing differences in underlying neurobiology in relation to empathy amongst those qualifying for the LPE specifier.

### Network Density was Different Between Groups

The LPE specifier group had fewer positive connections within the DMN for both cognitive and affective models. Similarly, within the SAL, the LPE specifier group had fewer positive connectivity for both cognitive and affective models. This supports prior work demonstrating less DMN [15, 17] and SAL [22] connectivity at higher CU traits and extends it to demonstrate this directly indicate differences in empathy amongst those in the clinically relevant LPE specifier.

Within the ECN, the LPE specifier group had more positive connections in the cognitive model. While initially this may seem to contradict prior work demonstrating less density in the ECN at higher CU traits [19] this prior finding came from work predicting CU traits rather than empathy in relation to the clinically relevant LPE specifier. Given that the LPE specifier group had substantially lower cognitive empathy and the normative group cognitive model did not generalize to them, the LPE specifier group could plausibly be over relying on top-down cognitive processes beyond their capacity to support cognitive empathy, which could underlie cognitive empathy impairments.

Additionally, we revealed unique patterns of fewer connections in the limbic network as well as fewer connections in the cerebellar system for both cognitive and affective models. The limbic system is involved in regulation of multiple systems and is also, related to the present study, largely involved in emotion responsiveness [47]. Less connections in this network in relation to both cognitive and affective empathy may indicate deficits in coordinating systems related to emotional responsiveness necessary to engage in empathy. Similarly, the cerebellar network has been implicated in social skills, attention, and empathy [27]. Less density in the cerebellar network plausibly make is more difficult for this network to support empathy evidenced by those in the LPE group that have substantially less empathy on average also have less connections in the cerebellar network. These findings suggest that it is important to consider subcortical networks to add further context of the brain’s associations with empathy amongst youth with CU traits and the LPE specifier.

The LPE specifier groups had more positive connections between ECN-DMN in the cognitive model and fewer negative connections in the affective model. This finding addresses a dichotomy in the literature in which one finding revealed more positive density between ECN-DMN at higher CU traits that was suggested to support cognitive empathy deficits [19, 46, 48] whereas another study found CU traits moderated the association between ECN-DMN connectivity and affective empathy [24]. Because we expected more negative connections and fewer positive connections between these networks in healthy populations [49] this inverse association from normative participants supports aberrant connectivity between these networks that are distinct for cognitive and affective empathy. This is consistent with the view that competition between ECN-DMN supports empathy generally [23] and extends it to demonstrate discrete impairments in support of empathy between normative and LPE specifier groups.

Inter-network SAL-DMN connectivity demonstrated both fewer positive and more negative connections in the cognitive model in addition to fewer positive connections in the affective model. Although prior work in this area has examined the magnitude of SAL-DMN connection [15], this work did not consider the relationships with empathy or the functional connection associations with empathy amongst those in the clinically relevant LPE specifier. Thus, the present finding is novel and, coupled with the finding of fewer positive connections between SAL-ECN for both cognitive and affective empathy, suggests that when engaging in empathy there may be a larger reliance on switching to top-down cognitive processes for empathy. There may be inadequate resources for these tasks resulting in impaired empathy.

These results must be interpreted under the following limitations. First, we assumed homogeneity amongst theoretically derived groups of those that are normative and those in the LPE specifier group. However, there is substantial evidence of heterogeneity amongst adolescents on the spectrum of CU traits [e.g., 19, 50]. Although we derived individual-level matrices for our analysis, future work would benefit from characterizing this heterogeneity in future work. Second, the sample size was rather modest (n=86) which may have missed some important effects. And finally, the sample was from the community which may not reflect forensic samples. Future work would benefit from including both community and forensic samples into the analysis and test if there are differences.

## Conclusion

The present study demonstrated unique patterns of functional connectivity predict empathy that are distinctly different between normative and the theoretically derived clinical LPE specifier. Further this work demonstrates density differences that partially support prior work and extend it to demonstrate density differences in relation to cognitive and affective empathy directly between normative and LPE specifier groups. This work adds to the existing literature demonstrating network differences in relation to empathy and CU traits that, with further work, may evidence a unique connectivity profile underlying empathy impairment in youth with CU traits. These findings may advance the understanding of empathic impairments amongst those with the LPE specifier in community samples and may have applications in diagnosing and intervention of severe CU traits.

## References

1. Frick, P.J., et al., Annual research review: A developmental psychopathology approach to understanding callous-unemotional traits in children and adolescents with serious conduct problems. Journal of child Psychology and Psychiatry, 2014. 55(6): p. 532–548.

2. Frick, P.J., et al., Can callous-unemotional traits enhance the understanding, diagnosis, and treatment of serious conduct problems in children and adolescents? A comprehensive review. Psychological Bulletin, 2014. 140(1): p. 1.

3. American Psychiatric Association, Diagnostic and Statistical Manual of Mental Disorders. 5th ed. 2013, Arlington, VA: American Psychiatric Publishing.

4. World Health Organization, International statistical classification of diseases and related health problems. 11th ed. 2020.

5. Frick, P.J. and S.F. White, Research review: The importance of callous-unemotional traits for developmental models of aggressive and antisocial behavior. Journal of child psychology and psychiatry, 2008. 49(4): p. 359–375.

6. Kimonis, E.R., et al., Using self-reported callous-unemotional traits to cross-nationally assess the DSM-5 ‘With Limited Prosocial Emotions’ specifier. Journal of Child Psychology and Psychiatry, 2015. 56(11): p. 1249–1261.

7. Rijnders, R.J., et al., Unzipping empathy in psychopathy: Empathy and facial affect processing in psychopaths. Neuroscience & Biobehavioral Reviews, 2021.

8. Decety, J., et al., Empathy as a driver of prosocial behaviour: Highly conserved neurobehavioural mechanisms across species. Philosophical Transactions of the Royal Society B: Biological Sciences, 2016. 371(1686): p. 20150077.

9. Decety, J., Dissecting the neural mechanisms mediating empathy. Emotion Review, 2011. 3(1): p. 92–108.

10. Eisenberg, N., N.D. Eggum, and L. Di Giunta, Empathy-related responding: Associations with prosocial behavior, aggression, and inter-group relations. Social issues and policy review, 2010. 4(1): p. 143–180.

11. Winters, D.E., W. Wu, and S. Fukui, Longitudinal Effects of Cognitive and Affective Empathy on Adolescent Substance Use. Substance Use & Misuse, 2020: p. 1–7.

12. Winters, D.E., et al., Systematic review and meta-analysis of socio-cognitive and socio-affective processes association with adolescent substance use. Drug and Alcohol Dependence, 2020: p. 108479.

13. Tompson, S.H., et al., Network approaches to understand individual differences in brain connectivity: opportunities for personality neuroscience. Personality Neuroscience, 2018. 1.

14. Winters, D.E., et al., Network functional connectivity underlying dissociable cognitive and affective components of empathy in adolescence. Neuropsychologia, 2021. 156: p. 107832.

15. Winters, D.E. and L.W. Hyde, Associated functional network connectivity between callous-unemotionality and cognitive and affective empathy. Journal of Affective Disorders, 2022.

16. Corbetta, M., G. Patel, and G.L. Shulman, The reorienting system of the human brain: from environment to theory of mind. Neuron, 2008. 58(3): p. 306–324.

17. Umbach, R.H. and N. Tottenham, Callous-unemotional traits and reduced default mode network connectivity within a community sample of children. Development and Psychopathology, 2021. 33(4): p. 1170–1183.

18. Winters, D.E., et al., Resting-state connectivity underlying cognitive control’s association with perspective taking in callous-unemotional traits. bioRxiv, 2022.

19. Winters, D.E., J.T. Sakai, and M.R. Carter, Resting-state network topology characterizing callous-unemotional traits in adolescence. NeuroImage: Clinical, 2021. 32: p. 102878.

20. Bilevicius, E., et al., Trait Emotional Empathy and Resting State Functional Connectivity in Default Mode, Salience, and Central Executive Networks. Brain Sci, 2018. 8(7).

21. Winters, D.E., et al., Cognitive and Affective Empathy as Indirect Paths Between Heterogeneous Depression Symptoms on Default Mode and Salience Network Connectivity in Adolescents. Child Psychiatry & Human Development, 2021.

22. Yoder, K.J., B.B. Lahey, and J. Decety, Callous traits in children with and without conduct problems predict reduced connectivity when viewing harm to others. Scientific Reports, 2016. 6(1): p. 20216.

23. Xin, F. and X. Lei, Competition between frontoparietal control and default networks supports social working memory and empathy. Social cognitive and affective neuroscience, 2015. 10(8): p. 1144–1152.

24. Winters, D.E., et al., Callous-unemotional traits in adolescents moderate neural network associations with empathy. Psychiatry Research: Neuroimaging, 2022. 320: p. 111429.

25. Ebisch, S.J.H., et al., Intrinsic Shapes of Empathy: Functional Brain Network Topology Encodes Intersubjective Experience and Awareness Traits. Brain Sci, 2022. 12(4).

26. Hamada, M., et al., People with High Empathy Show Increased Cortical Activity around the Left Medial Parieto-Occipital Sulcus after Watching Social Interaction of On-Screen Characters. Cerebral Cortex, 2022.

27. Hoche, F., et al., Cerebellar Contribution to Social Cognition. Cerebellum, 2016. 15(6): p. 732–743.

28. Finn, E.S., et al., Functional connectome fingerprinting: identifying individuals using patterns of brain connectivity. Nature neuroscience, 2015. 18(11): p. 1664–1671.

29. Shen, X., et al., Using connectome-based predictive modeling to predict individual behavior from brain connectivity. nature protocols, 2017. 12(3): p. 506–518.

30. Ye, S., et al., Connectome-based model predicts individual psychopathic traits in college students. Neuroscience Letters, 2022. 769: p. 136387.

31. Yao, G., et al., Neural mechanisms underlying empathy during alcohol abstinence: evidence from connectome-based predictive modeling. Brain Imaging and Behavior, 2022: p. 1–10.

32. Wechsler, D., Wechsler abbreviated scale of intelligence-(WASI-II). Vol. 4. 2011, San Antonio, TX: NCS Pearson.

33. Nooner, K.B., et al., The NKI-Rockland Sample: A Model for Accelerating the Pace of Discovery Science in Psychiatry. Frontiers in Neuroscience, 2012. 6(152).

34. Achenbach, T.M., Manual for the youth self-report and 1991 profile. 1991: Department of Psychiatry, University of Vermont Burlington, VT.

35. Petersen, A.C., et al., A self-report measure of pubertal status: Reliability, validity, and initial norms. Journal of Youth and Adolescence, 1988. 17(2): p. 117–133.

36. Whitfield-Gabrieli, S. and A. Nieto-Castanon, Conn: a functional connectivity toolbox for correlated and anticorrelated brain networks. Brain Connect, 2012. 2(3): p. 125–41.

37. Penny, W.D., et al., Statistical parametric mapping: the analysis of functional brain images. 2011: Elsevier.

38. Glasser, M.F., et al., The minimal preprocessing pipelines for the Human Connectome Project. Neuroimage, 2013. 80: p. 105–124.

39. Wu, C.W., et al., Empirical evaluations of slice-timing, smoothing, and normalization effects in seed-based, resting-state functional magnetic resonance imaging analyses. Brain connectivity, 2011. 1(5): p. 401–410.

40. Desikan, R.S., et al., An automated labeling system for subdividing the human cerebral cortex on MRI scans into gyral based regions of interest. Neuroimage, 2006. 31(3): p. 968–980.

41. Tzourio-Mazoyer, N., et al., Automated anatomical labeling of activations in SPM using a macroscopic anatomical parcellation of the MNI MRI single-subject brain. Neuroimage, 2002. 15(1): p. 273–289.

42. Pedregosa, F., et al., Scikit-learn: Machine learning in Python. the Journal of machine Learning research, 2011. 12: p. 2825–2830.

43. Ju, Y., et al., Connectome-based models can predict early symptom improvement in major depressive disorder. Journal of affective disorders, 2020. 273: p. 442–452.

44. Varoquaux, G., et al., Brain covariance selection: better individual functional connectivity models using population prior. Advances in neural information processing systems, 2010. 23.

45. Dosenbach, N.U., et al., Prediction of individual brain maturity using fMRI. Science, 2010. 329(5997): p. 1358–1361.

46. Pu, W., et al., Alterations of Brain Functional Architecture Associated with Psychopathic Traits in Male Adolescents with Conduct Disorder. Sci Rep, 2017. 7(1): p. 11349.

47. Isaacson, R., The limbic system. 2013: Springer Science & Business Media.

48. Dotterer, H.L., et al., Connections that characterize callousness: Affective features of psychopathy are associated with personalized patterns of resting-state network connectivity. NeuroImage: Clinical, 2020. 28: p. 102402.

49. Uddin, L.Q., et al., Functional connectivity of default mode network components: correlation, anticorrelation, and causality. Human brain mapping, 2009. 30(2): p. 625–637.

50. Winters, D.E., et al., Efficiency of heterogenous functional connectomes explains variance in callous-unemotional traits after computational lesioning of cortical midline and salience regions. bioRxiv, 2022: p. 2022.10.07.511379.

